# Assessing protein quality, reactive lysine and gastric digestion characteristics of feed-grade black soldier fly meal: Implications for post-weaning piglet feed formulation

**DOI:** 10.64898/2026.08.19.745673

**Authors:** Hugo Luttenschlager, Niels Wever, Samuel Jansseune, Wouter Hendriks, Yves Beckers, Joachim Carpentier, Aurore Richel, Frédéric Francis, Rudy Caparros Megido

## Abstract

Insect proteins represent a promising and more sustainable alternative to soybean meal in swine nutrition. However, high dietary inclusion levels often result in reduced animal performance, indicating that factors other than crude protein concentration should be considered. This study evaluated the protein quality and degradability of feed-grade defatted black soldier fly (BSF) meal in comparison with a conventional organic protein core for post-weaning piglets. Protein content, amino acid composition, reactive lysine, Maillard reaction markers, buffering capacity, and protein hydrolysability were determined.

Our results showed that defatted BSF meal exhibited a protein content and amino acid profile comparable to those of the protein core but contained substantially higher levels of free amino acids. Accounting for free amino acids reduced the estimated lysine damage from 18.41% to 10.87%, highlighting the importance of considering free amino acids when determining reactive lysine. In addition, BSF meal exhibited a higher buffering capacity and lower protein degradability than the protein core.

These findings provide new insights into the nutritional limitations of feed-grade BSF meal and suggest that a more accurate assessment of reactive lysine, together with dietary strategies to reduce buffering capacity, may improve its utilization in post-weaning piglet diets.

## Introduction

Many countries rely on traditional protein sources for animal feed such as soybean meal (SBM) and fish meal (FM) (Jerzak & Śmiglak-Krajewska, 2020; Ndebele-Murisa et al., 2024; Wu et al., 2020). These resources, in addition to having production unevenly distributed relative to their consumption, raise several sustainability concerns, including deforestation, competition for arable land, and pressure on marine resources (Glencross et al., 2024; Taherzadeh & Caro, 2019). As a result, new locally produced protein sources have begun to emerge.

Among these new protein sources, insects have received considerable attention in recent years. This can be explained by their ease of rearing, rapid reproduction, as well as their high protein content and amino acid profile, allowing them to partially substitute conventional protein sources in various animal models, including monogastric and aquaculture species (Caparros Megido et al., 2024; da-Silva et al., 2024; Huis et al., 2020; Luttenschlager et al., 2026a). Among these insects, *Hermetia illucens* (L., 1758), also known as the black soldier fly (BSF), has attracted the interest of many farmers and researchers. This interest is driven by the fact that the larvae are polyphagous and voracious, the adults are highly prolific, and overall their production is relatively easy to implement at different scales (Hoc et al., 2019; Purkayastha & Sarkar, 2022). BSF therefore represents a valuable protein source capable of valorizing a wide range of organic materials such as unsold products, agricultural by-products, and even certain waste streams depending on current regulations (Chia et al., 2020; Liu et al., 2022; Suckling et al., 2021). Together, these characteristics make BSF an attractive ingredient for sustainable animal nutrition, particularly because it is produced for feed rather than food, thereby reducing feed-food competition.

However, within the European Union, these new protein sources fall under the category of processed animal proteins. Their production must therefore comply with specific regulations, including thermal processing steps aimed at reducing the risk of pathogen development or transmission as defined by Regulation (EC) No. 1069/2009 and its implementing Regulation (EU) No. 142/2011 (European Parliament and Council of the European Union, 2009, 2011). When these processed animal proteins undergo thermal treatments, particularly when prolonged or applied at high temperatures, there is a risk of altering their nutritional properties. Lysine, one of the first limiting amino acids, is particularly susceptible to heat-induced chemical modification because of the high reactivity of its ε-amino group (ε-NH₂) (Rutherfurd, 2015). These thermal treatments can reduce the proportion of reactive lysine in feed ingredients, mainly through reactions involving the ε-amino group with other reactive compounds. Among these, Maillard reactions, involving reducing sugars, represent a major pathway for lysine modification. If the damage is further exacerbated, this will lead to Amadori products and then irreversibly to advanced glycation end products (AGEs) (Teodorowicz et al., 2018; Twarda-Clapa et al., 2022; Wei et al., 2018). These reactions reduce lysine availability, rendering it metabolically unavailable and potentially leading to decreased zootechnical performance (Deng et al., 2017; Loveday, 2023; Zenker et al., 2020). It is therefore important to evaluate lysine availability and the presence of Maillard reaction markers in novel protein sources subjected to thermal processing, to refine diet formulation and better meet amino acid requirements.

To our knowledge, this study is the first to compare the amino acid composition and protein quality of a protein core from a standard organic piglet feed with a defatted black soldier fly meal of feed-grade quality. This research specifically aims to evaluate lysine availability by distinguishing total lysine from reactive lysine, and to assess the contribution of Maillard reactions, including both early and advanced products, to the loss of available lysine. In addition, the analysis of free amino acids, total amino acid profiles, buffering capacity, and protein degrees of hydrolysis will provide further insight into protein quality and potential analytical biases. Finally, these analyses will help determine the extent to which these protein sources complement the corresponding energy core in terms of amino acid balance and effective lysine availability, to improve feed formulation strategies.

## Materials and methods

The control protein core is a PRODABIO (Awans, Belgium) post-weaning piglet protein concentrate that has already been used and described in a previous experiment (Luttenschlager et al., 2026b). We also analyzed the energy core from the same supplier to determine the extent to which the energy core complements the protein core or the BSF meal.

### BSF meal production

The production of BSF-derived materials complied with Belgian and European regulations for animal feed. The processing of BSF meal was conducted under official authorizations granted by the competent authorities (Federal Agency for the Safety of the Food Chain and Walloon Region), including approval for insect producers other than bees and bumblebees, handling category 3 animal by-products and the manufacture of compound feeds containing processed animal proteins derived from insects.

BSF larvae were produced under controlled rearing conditions (25.3 °C ± 0.48°C and 57.50% ± 14.68% humidity) and harvested before pupation at the functional and evolutionary entomology laboratory Gembloux Agro-Bio Tech (University of Liege). After harvesting, larvae were frozen at - 20°C, crushed (Steel Fruit Crusher 1100 W, WilTec, Eschweiler, Germany), and oven-dried at 70°C for 8 h.

Lipids were subsequently reduced to 5.56 % of the dried larval meal at Extratex S.F.I. (Pont-Saint-Vincent, France) using supercritical CO₂ extraction under industrial conditions (400 L, 60 °C, 600 bar, 5 h), yielding a defatted BSF meal as described in a previous work (Luttenschlager et al., 2026b).

The defatted BSF was then subjected to a thermal treatment consisting of 80 °C for 120 min followed by 100 °C for 60 min, in accordance with European regulations for processed animal proteins Regulation (EC) No. 1069/2009 and its implementing Regulation (EU) No. 142/2011 (European Parliament and Council of the European Union, 2009, 2011). The resulting BSF meal was finally ground before further analyses with a hammer mill (SM 100, Retsch GmbH, Germany) equipped with a 3 mm sieve.

### Sample weighing and preparation

All samples were weighed using a precision balance (Mettler Toledo XP205 analytical balance, Greifensee, Switzerland). Samples were ground using (20 Hz, 5min, agate balls and containers, MM200, Retsch, Haan, Germany), except for free amino acid and tryptophan analyses, for which the finely ground meals were used as such.

### Nitrogen content

Nitrogen content was determined using the Dumas method (AOAC 990.03) and used to calculate crude protein content (Wendt Thiex & Latimer, 2023).

### Total amino acid analysis

Total and free amino acid analyses were performed in accordance with ISO 13903 and ISO 13904 (ISO, 2005a, 2005b).

### Performic oxidation for sulfur amino acids

Samples were oxidized for 16 h at 0 °C in 0.1 mL of a 9:1 (v/v) formic acid-hydrogen peroxide solution in sealed tubes. Subsequently, samples were brought to room temperature and 16.8 mg of sodium disulfite was added to each sample.

### Acid hydrolysis for amino acids

In each tube, 1 mL of a hydrolysis mixture was added to each sample (1 g phenol dissolved in 200 mL water, followed by gradual addition of 585 g of concentrated hydrochloric acid and dilution to 1 L with water).

### Amino acid analysis

After performic oxidation of sulfur amino acids and preparation of non-oxidized samples, all samples were placed under vacuum and flame-sealed in glass tubes. Samples were then hydrolyzed for 23 h at 110 °C. The glass tubes were subsequently opened using a diamond-tipped pen to score the surface of the tube, after which a small drop of water and molten glass was applied to the score. 0.5 µmol of norleucine was added to each tube, and the solvants were subsequently evaporated using a SpeedVac concentrator (Savant SC250EXP, Savant Instruments Inc., Farmingdale, NY).

For sulfur amino acids (oxidized samples), samples were dried for 1 h 45 min at 45 °C until nearly dry. Subsequently, 2 mL of sample loading buffer was added, followed by adjustment of the pH to 2.2 ± 0.2.

For non-sulfur amino acids (non-oxidized samples), samples were subjected to a two-step heating program consisting of 30 min at 45 °C followed by 2 h at 75 °C until dryness. Subsequently, 2 mL of sample dilution buffer was added to each tube.

### Free amino acid analysis

One gram of sample was weighed into a 50 mL polypropylene tube. Then, 20 mL of 0.1 M HCl was added and free amino acids were extracted by horizontal shaking for 30 min. After centrifugation at 3000 × g for 10 min, 10 mL of supernatant was mixed with 5 mL of 6% sulfosalicylic acid. After 10 min, the tubes were centrifuged again at 3000 × g and 9.5 mL of supernatant was mixed with 0.5 mL of norleucine to obtain a final concentration of 0.25 mM. Finally, the pH was adjusted to 2.20 by addition of a 4 M sodium hydroxide solution. Cation-exchange analysis

All samples were then mixed and filtered over 0.2 µm nylon syringe filters before analysis. Amino acids were then analyzed by ion-exchange chromatography using a Sykam S 5200 cation-exchange high-performance liquid chromatography (HPLC) system coupled with a Sykam S 7130 post-column derivatization unit (Sykam, Eresing, Germany). Separation was achieved using a LCAK04/Na precolumn (100 × 4.6 mm) and a LCAK13/Na analytical column (175 × 4.6 mm) and sodium buffers. Following post-column derivatization with ninhydrin, amino acids were detected photometrically at 570 nm, except for proline, which was detected at 440 nm.

### Alkaline hydrolysis for tryptophan

Tryptophan was determined after alkaline hydrolysis with barium hydroxide octahydrate at 110 °C for 23 h. After cooling, α-methyltryptophan was added as an internal standard, and the hydrolysates were neutralized and adjusted to pH 3 before dilution to volume with methanol and subsequent chromatographic analysis using reversed-phase chromatography and fluorescence detection.

### Reactive lysine analysis (OMIU-lysine)

The preparation of the 0.6 M OMIU solution was adapted from the literature (Hulshof et al., 2017; Moughan C Rutherfurd, 1996). For that, 8.4 g of barium hydroxide octahydrate was added to approximately 32 ml of CO2-free (boiled for 10 min) distilled deionized water. Then, 4 g of O-methylisourea was added to the centrifuge tube. Once at room temperature, the solution was centrifuged at 6400 × g for 10 min at 20°C using a high-speed centrifuge (Beckman Colter, California, USA). The supernatant was collected, and the remaining pellet was washed with approximately 2 mL of boiled distilled deionized water, followed by a second centrifugation step.

The resulting supernatants were pooled, and the pH was checked to confirm it exceeded 12. The pH was then adjusted to 11 using 6 M HCl, and the volume was brought to 40 mL with boiled distilled deionized water. In each glass tube containing the samples and controls (peas, rapeseed meal, faba beans, L-lysine, and the dipeptide Lys–Lys), 1 mL of 0.6 M OMIU was added. The tubes were sealed and placed on a shaker for 7 days. Samples were dried under vacuum using a SpeedVac concentrator, then hydrolyzed using hydrochloric acid, reevaporated, and reconstituted as described for the analysis of total amino acids. Analysis of homo-arginine was performed using cation-exchange chromatography using sodium buffers with the use of a divert-valve to divert the ammonia peak before the elution of arginine and homo-arginine.

### Advanced Glycation End-product Analysis

The following advanced glycation end products were analyzed: Nε-(carboxymethyl)lysine (CML), Nε-(carboxyethyl)lysine (CEL), and methylglyoxal-derived hydroimidazolone (MG-H1). MG-H1 is not a lysine marker but was also analyzed as an arginine-derived marker of heat-induced protein damage. The protocol was adapted from the current literature (Scheijen et al., 2016).

In each tube containing 10 mg of samples, 60 µL of 0.2 M borate buffer (pH 9.2) was added before 40 µL of reduction reagent containing 1 M sodium borohydride in 0.1 M sodium hydroxide. After 4 h of incubation, 0.9 mL of hydrolysis mixture was added to each tube, which was then vacuum-sealed as described previously and placed in an oven at 110 °C for 23 h.

The glass tubes were reopened as described above, after which 500 µL of an internal standard solution containing CEL-d4, CML-d4, MG-H1-d4 and furosine-d4 diluted in 50% acetonitrile was added to each tube. The samples were dried in a SpeedVac at 45 °C and the residue was reconstituted in 50% mobile phase A (20 mM ammonium formate, pH 3) and B (0.1% formic acid in acetonitrile) before analysis. Finally, the samples were filtered through a 0.22 µm nylon filter into 1.5 mL glass vials before analysis.

Furosine was analyzed using the same procedure as described above, omitting the reduction step.

Analysis was performed using liquid chromatography mass spectrometry (LCMS) on an Aquity LC coupled to a Quattro Premier XE triple quadrupole mass spectrometer (Waters, Milford, US). Separation was performed using hydrophilic interaction liquid chromatography on an Accucore HILIC (100 x 2.1mm, 2.6 µM) (Thermo, Waltham, US). The column was maintained at 50°C, and the initial condition was 30% mobile phase A, which was held for 1 minute, after which a linear ramp up to 60% A for 6 minutes was performed. This was then held for 2 minutes before returning to the initial conditions and subsequent equilibration of 4 minutes. Ionization was achieved using electrospray ionization at 3 kV and 650 L/min spray flow. Ions were measured in MRM using argon as collision gas and were optimized for the system by infusion of a standard solution (Table S1); the dwell time was adjusted for each component to provide at least 15 measurements per peak.

### Buffer capacity and pH-stat analysis

The buffering capacity was calculated based on literature (Lawlor et al., 2005) with a few modifications. Samples were analyzed in duplicate. An amount of 500 mg of ball ground sample was weighed into a 20 mL glass vial (VWR, Leuven, Belgium). Then, 10 mL of 0.01 M HCl was added to the sample together with a magnetic stir bar, and the mixture was stirred (500rpm) at 4°C overnight. The vial was heated for 30 min in a water bath at 37°C and then transferred to a thermostated jacketed beaker maintained at 39°C and fitted with a lid. The pH was automatically decreased to pH 2 using 0.01 M HCl in 0.5 pH-unit steps whenever the pH remained stable for 360s. The volume of HCl required to decrease the pH from its initial value to pH 2 was recorded and used as a measure of the buffering capacity of the sample.

The pH-stats hydrolysis by porcine pepsin were based on an adaptation from the literature (Mat et al., 2018). Samples were ball ground as escribed for buffering capacity determination. Then, 100 mg of protein, based on total amino acid content, were weighed into 20 mL glass vials. The pH of each sample was decreased to 2.5 with 0.5 M HCl and the volume was brought to 10 mL with deionized water. The amount of 0.5 M HCl added was determined according to the buffering capacity results to reach a pH of 2.5 after equilibration. Water was added first to prevent a too high acid concentration in contact with the samples. Samples were stirred (500rpm) at 4°C overnight. Each vial was then heated for 30 min in a water bath at 37°C and subsequently transferred to a thermostated jacketed beaker maintained at 39°C and fitted with a lid. The pH was manually adjusted to approximately 2.510 using concentrated HCl and NaOH and then automatically adjusted to pH 2.500 using 0.1M HCl by the titration unit (Titrando 902, Methrom, Herisau, CH). Afterward, 125 µL of a 4 mg/mL pepsin (P7012, Sigma, Burlington, US) solution at pH 2.5 was added. During the 7h hydrolysis, the pH was continuously monitored using a pH electrode connected to an automatic titration system, which maintained the pH at 2.5 by the addition of 0.1 M HCl. Protein hydrolysability was also assessed using the previously described pH-stat method, except that hydrolysis was carried out for 12 h using 100yl of a 60 mg/ml pepsin solution at pH 2.5. This high pepsin dose combined with a long hydrolysis time was used to indicates the maximal potential DH that could be reached while the low pepsin dose was used to have more resolution in differentiating the hydrolysis rate.

The cumulative volume of titrant added to maintain a constant pH was continuously recorded and used to calculate the degree of hydrolysis (DH). At a pH of 2.5, DH can be calculated using the following equation:

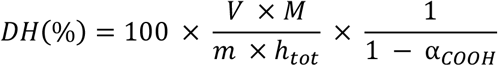

where DH is the degree of protein hydrolysis, V is the volume of titrant (ml), M is the normality of the titrant (meq/ml), and *m* is the mass of protein (g). *h*tot is the number of peptide bonds per gram of protein. The total hydrolysable peptide bond content (H_tot_) was estimated from the amino acid composition of each ingredient. The molar amount of each amino acid was calculated from its total amino acid content and molecular weight. To account for water loss during peptide bond formation, the residue molecular weight (molecular weight − 18.01 g mol^−1^) was used. H_tot_ was expressed as mmol of hydrolysable peptide bonds per gram of protein.*αCOOH* is the mean degree of dissociation of the carboxylic groups, and was estimated to be 0.075 at pH 2.5 and 37 °C (Mat et al., 2018; Yasumaru & Lemos, 2014).

### Mineral analysis

Minerals were measured using inductively coupled plasma optical emission spectrometry (ICO-OES) after microwave digestion (MARS6, CEM, Matthews, US). In short, 0.5 grams of sample material was weighed into a Teflon digestion vessel, 8 ml of nitric acid and 2 ml of hydrochloric acid were added, and the tubes were left open in a fume hood for 30 minutes after which they were closed. The tubes were then placed in the microwave and ramped to 200°C over 20 minutes and held at that temperature for a final 20 minutes. The content of the tubes were then quantitatively rinsed into 100 ml volumetric flasks with distilled water. A final dilution was made using Scandium at a final concentration of 5 mg/L as internal standard to correct for matrix effects. Measurements were conducted against external calibration. For all minerals but Sodium and Potassium two emission bands were chosen based on the manufacturer’s recommendations, to determine if there was spectral interference during the measurement.

## Results

### Amino acid composition and protein estimation

The Dumas analysis showed that the energy core contained 18.3 ± 0.6 g/kg DM of nitrogen and that the BSF meal contained more nitrogen (98.7 ± 0.3 g/kg DM) than the protein core (85.7 ± 0.3 g/kg DM). Using a nitrogen-to-protein conversion factor of 6.25, this corresponds to a crude protein content of 114.3 ± 3.8 g/kg DM for the energy core, 535.5 ± 1.6 g/kg DM for the protein core, and 616.9 ± 1.8 g/kg DM. For BSF meal, a conversion factor of 4.76 has been recommended to avoid overestimating protein content (Janssen et al., 2017), resulting in an estimated protein content of 469.8 ± 1.4 g/kg DM.

Our amino acid analysis (Table 1) showed that the total amino acids contents of the energy core, protein core, and BSF meal were 102.0 ± 1.7 g/kg DM, 497.4 g/kg ± 6.7 DM, and 499.8 ± 6.0 g/kg DM, respectively. BSF meal also distinguished itself by its higher free amino acid content, reaching 56.2 ± 0.7 g/kg DM, which represented 11.2% of total amino acids. This was substantially higher than in the protein core (3.0 ± 0.3 g/kg DM; 0.6% of total amino acids) and the energy core (2.2 ± 1.6 g/kg DM; 2.2% of total amino acids).

Glucosamine, used as a proxy for chitin-derived compounds, was detected at concentrations of 0 g/kg DM in the energy core, 0.8 g/kg DM in the protein core, and 31.2 g/kg DM in the BSF meal.

**Table 1:** Free and total amino acid profiles in g/kg of the energy core, protein core, and defatted BSF meal. Sum AA = Total amino acids; Sum IAA = Total indispensable amino acids; Sum NIAA = Total non-indispensable amino acids; IAA/NIAA = Ratio of indispensable amino acids to non-indispensable amino acids

| Feed | BSF Meal | BSF Meal | Protein Core | Protein Core | Energy Core | Energy Core |
| --- | --- | --- | --- | --- | --- | --- |
| Form | Total | Free | Total | Free | Total | Free |
| Unit | g/kg DM | g/kg DM | g/kg DM | g/kg DM | g/kg DM | g/kg DM |
| Alanine | $39.7 \pm 0.2$ | $9.1 \pm 0.1$ | $23.0 \pm 0.2$ | $0.3 \pm 0.0$ | $5.2 \pm 0.1$ | $0.1 \pm 0.0$ |
| Arginine | $24.0 \pm 0.1$ | $2.1 \pm 0.0$ | $34.0 \pm 0.4$ | $1.0 \pm 0.1$ | $6.2 \pm 0.1$ | $0.1 \pm 0.0$ |
| Aspartic acid | $46.9 \pm 0.2$ | $2.7 \pm 0.0$ | $57.1 \pm 0.4$ | $0.4 \pm 0.1$ | $7.6 \pm 0.2$ | $0.1 \pm 0.0$ |
| Cysteine | $4.1 \pm 0.1$ | $0.0 \pm 0.0$ | $7.3 \pm 0.4$ | $0.0 \pm 0.0$ | $2.4 \pm 0.0$ | $0.0 \pm 0.0$ |
| Glutamic acid | $57.1 \pm 1.7$ | $8.1 \pm 0.1$ | $76.4 \pm 0.6$ | $0.6 \pm 0.1$ | $22.0 \pm 0.3$ | $0.2 \pm 0.0$ |
| Glycine | $30.1 \pm 0.2$ | $2.1 \pm 0.0$ | $22.5 \pm 0.2$ | $0.0 \pm 0.0$ | $4.9 \pm 0.1$ | $0.0 \pm 0.0$ |
| Histidine | $17.7 \pm 0.6$ | $2.4 \pm 0.0$ | $13.2 \pm 0.1$ | $0.2 \pm 0.0$ | $2.7 \pm 0.0$ | $1.1 \pm 1.5$ |
| Isoleucine | $24.1 \pm 0.3$ | $2.9 \pm 0.0$ | $23.8 \pm 0.4$ | $0.0 \pm 0.0$ | $3.9 \pm 0.1$ | $0.0 \pm 0.0$ |
| Leucine | $37.7 \pm 0.2$ | $4.2 \pm 0.0$ | $42.0 \pm 0.6$ | $0.0 \pm 0.0$ | $8.0 \pm 0.2$ | $0.0 \pm 0.0$ |
| Lysine | $30.1 \pm 0.1$ | $3.1 \pm 0.0$ | $33.5 \pm 0.5$ | $0.1 \pm 0.0$ | $4.4 \pm 0.1$ | $0.1 \pm 0.0$ |
| Methionine | $8.6 \pm 0.2$ | $0.5 \pm 0.0$ | $8.1 \pm 0.6$ | $0.0 \pm 0.0$ | $1.9 \pm 0.1$ | $0.0 \pm 0.0$ |
| Phenylalanine | $23.4 \pm 0.1$ | $1.9 \pm 0.0$ | $27.6 \pm 0.3$ | $0.1 \pm 0.0$ | $5.0 \pm 0.1$ | $0.0 \pm 0.0$ |
| Proline | $34.2 \pm 1.0$ | $6.0 \pm 0.4$ | $26.1 \pm 0.4$ | $0.0 \pm 0.0$ | $8.8 \pm 0.1$ | $0.1 \pm 0.0$ |
| **Serine** | 22.8 ± 0.2 | 2.2 ± 0.0 | 25.7 ± 0.4 | 0.0 ± 0.0 | 4.7 ± 0.1 | 0.0 ± 0.0 |
| **Threonine** | 22.4 ± 0.2 | 3.6 ± 0.1 | 23.0 ± 0.3 | 0.1 ± 0.1 | 3.9 ± 0.0 | 0.0 ± 0.0 |
| **Tryptophane** | 8.7 ± 0.1 | 0.0 ± 0.0 | 6.1 ± 0.4 | 0.0 ± 0.0 | 1.3 ± 0.0 | 0.0 ± 0.0 |
| **Tyrosine** | 33.9 ± 0.3 | 1.4 ± 0.0 | 21.7 ± 0.4 | 0.1 ± 0.0 | 3.9 ± 0.0 | 0.0 ± 0.0 |
| **Valine** | 34.5 ± 0.4 | 4.1 ± 0.0 | 26.5 ± 0.3 | 0.1 ± 0.0 | 5.2 ± 0.1 | 0.0 ± 0.0 |
| **Sum AA** | 499.8 ± 6.0 | 56.2 ± 0.7 | 497.4 ± 6.7 | 3.0 ± 0.3 | 102.0 ± 1.7 | 2.2 ± 1.6 |
| **Sum IAA** | 231.0 ± 2.3 | 24.6 ± 0.2 | 237.7 ± 3.9 | 1.5 ± 0.2 | 42.6 ± 0.9 | 1.4 ± 1.5 |
| **Sum NIAA** | 268.8 ± 3.7 | 31.6 ± 0.5 | 259.8 ± 2.8 | 1.4 ± 0.1 | 59.3 ± 0.8 | 0.7 ± 0.1 |
| **IAA/NIAA** | 0.86 | 0.78 | 0.91 | 1.12 | 0.72 | 1.91 |

### Protein quality and Maillard markers

The OMIU guanidination assay revealed different levels of apparent lysine damage among the tested ingredients. Based on the initial measurements, the BSF meal exhibited 18.4% lysine damage (30.1 g/kg DM total lysine and 24.6 g/kg DM reactive lysine). In contrast, the protein core contained only 1.3% damage (33.5 g/kg DM total lysine and 33.1 g/kg DM reactive lysine). The energy core presented an intermediate level of damage of 14% (4.4 g/kg DM total lysine and 3.8 g/kg DM reactive lysine).

Additional experiments showed that the efficiency of OMIU-mediated guanidination differed according to the chemical form of lysine. The conversion yield was approximately 82% for the Lys-Lys dipeptide but only 27% for free L-lysine.

The BSF meal contained 3.1 g/kg DM of free lysine. Applying a correction based on the experimentally determined 27% conversion yield for free lysine increased the estimated reactive lysine content from 24.6 to 27.0 g/kg DM. It reduced the estimated lysine damage from 18.4% to 10.4%.

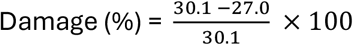

This damage to lysine can be partially detected through early Maillard markers (furosine) as well as certain AGEs. In addition, MG-H1, which is produced by the degradation of arginine, provides further evidence of the thermal origin of the damage to amino acids (Table 2).

**Table 2:** Composition of markers for early Maillard reactions, furosin (FUR), and advanced glycation end products (AGEs): Nε-(carboxymethyl)lysine (CML), Nε-(carboxyethyl)lysine (CEL), and methylglyoxal-derived hydroimidazolone (MG-H1) in BSF meal, the protein core, the energy core, and the control diet. Lysine (Lys) and reactive lysine (R-Lys) are included for comparison purposes.

| **Maillard markers** | **CML** | **CEL** | **FUR** | **MG-H1** | **R-Lys** | **Lys** |
| --- | --- | --- | --- | --- | --- | --- |
| **Unit** | mg/kg | mg/kg | mg/kg | mg/kg | mg/kg | mg/kg |
| **BSF meal** | 433.7 ± 42.3 | 459.5 ± 91.3 | 1545.1 ± 87.2 | 80.7 ± 9.4 | 27 ± 0.2 | 30.1 ± 0.1 |
| **Protein core** | 50.1 ± 18.0 | 54.6 ± 5.0 | 330.5 ± 30.6 | 101.5 ± 19.7 | 33.1 ± 2.4 | 33.5 ± 0.5 |
| **Energy core** | 7.5 ± 2.6 | 3.53 ± 1.0 | 64.2 ± 8.5 | 9.8 ± 3.6 | 3.8 ± 0.3 | 4.4 ± 0.1 |

### Mineral composition, buffering capacity, and protein degrees of hydrolysis

Selected mineral concentrations determined in the tested feeds are presented in Table 3. The total content of these minerals is 60.8 ± 1.0 g/kg DM for the BSF meal, 32.0 ± 0.7 g/kg DM for the protein core, and 33.1 ± 3.2 g/kg DM for the energy core.

**Table 3:** Mineral composition of BSF meal, protein core and energy core: Ca, P, Mg, Na, K, Fe, Mn, Zn, Co.

| Feed | BSF meal | Protein core | Energy core |
| --- | --- | --- | --- |
| **Ca g/kg** | $24.1 \pm 0.5$ | $2.7 \pm 0.1$ | $15.2 \pm 1.4$ |
| **P g/kg** | $10.6 \pm 0.1$ | $6.8 \pm 0.0$ | $6.4 \pm 0.1$ |
| **Mg g/kg** | $4.2 \pm 0.0$ | $2.3 \pm 0.1$ | $2.7 \pm 0.1$ |
| **Na g/kg** | $2.2 \pm 0.0$ | $0.6 \pm 0.0$ | $2.2 \pm 0.1$ |
| **K g/kg** | $18.3 \pm 0.2$ | $19.4 \pm 0.3$ | $5.9 \pm 0.1$ |
| **Fe mg/kg** | $411.4 \pm 15.6$ | $188.1 \pm 9.7$ | $337.2 \pm 37.0$ |
| **Mn mg/kg** | $176.3 \pm 3.6$ | $39.6 \pm 1.5$ | $206.7 \pm 43.4$ |
| **Zn mg/kg** | $817.2 \pm 32.0$ | $52.4 \pm 4.7$ | $187.8 \pm 69.6$ |
| **Co mg/kg** | $< 10.0$ | $< 10.0$ | $< 10.0$ |
| **Sum g/kg** | $60.8 \pm 1.0$ | $32.0 \pm 0.7$ | $33.1 \pm 3.2$ |

The buffering-capacity profiles differed among the test materials (Figure 1A). Across the pH intervals common to all three materials, BSF consistently showed the highest buffering capacity, increasing from 437.7 ± 30.3 mEq HCl kg⁻¹ DM pH⁻¹ between pH 4.5 and 4.0 to 1296.7 ± 50.3 mEq HCl kg⁻¹ DM pH⁻¹ between pH 2.5 and 2.0. The Protein core showed intermediate values, ranging from 297.0 ± 4.9 mEq HCl kg⁻¹ DM pH⁻¹ between pH 4.5 and 4.0 to 698.4 ± 9.8 mEq HCl kg⁻¹ DM pH⁻¹ between pH 2.5 and 2.0. The Energy core had a buffering capacity of 330.8 ± 49.1 mEq HCl kg⁻¹ DM pH⁻¹ between pH 4.5 and 4.0 and generally showed the lowest values below pH 4, reaching 572.4 ± 25.3 mEq HCl kg⁻¹ DM pH⁻¹ between pH 2.5 and 2.0. The cumulative acid-binding capacity profiles began at different initial pH values (Figure 1B). BSF had the highest initial pH, with a mean value of 6.8 ± 0.1, followed by the Energy core at 5.6 ± 0.0 and the Protein core at 4.9± 0.0. At pH 2, the cumulative acid-binding capacity (ABC-2) was 2881.7 ± 107.4 mEq HCl kg⁻¹ DM for BSF, 1143.9 ± 0.9 mEq HCl kg⁻¹ DM for the Protein core, and 1117.9 ± 15.8 mEq HCl kg⁻¹ DM for the Energy core.

**Figure 1.**
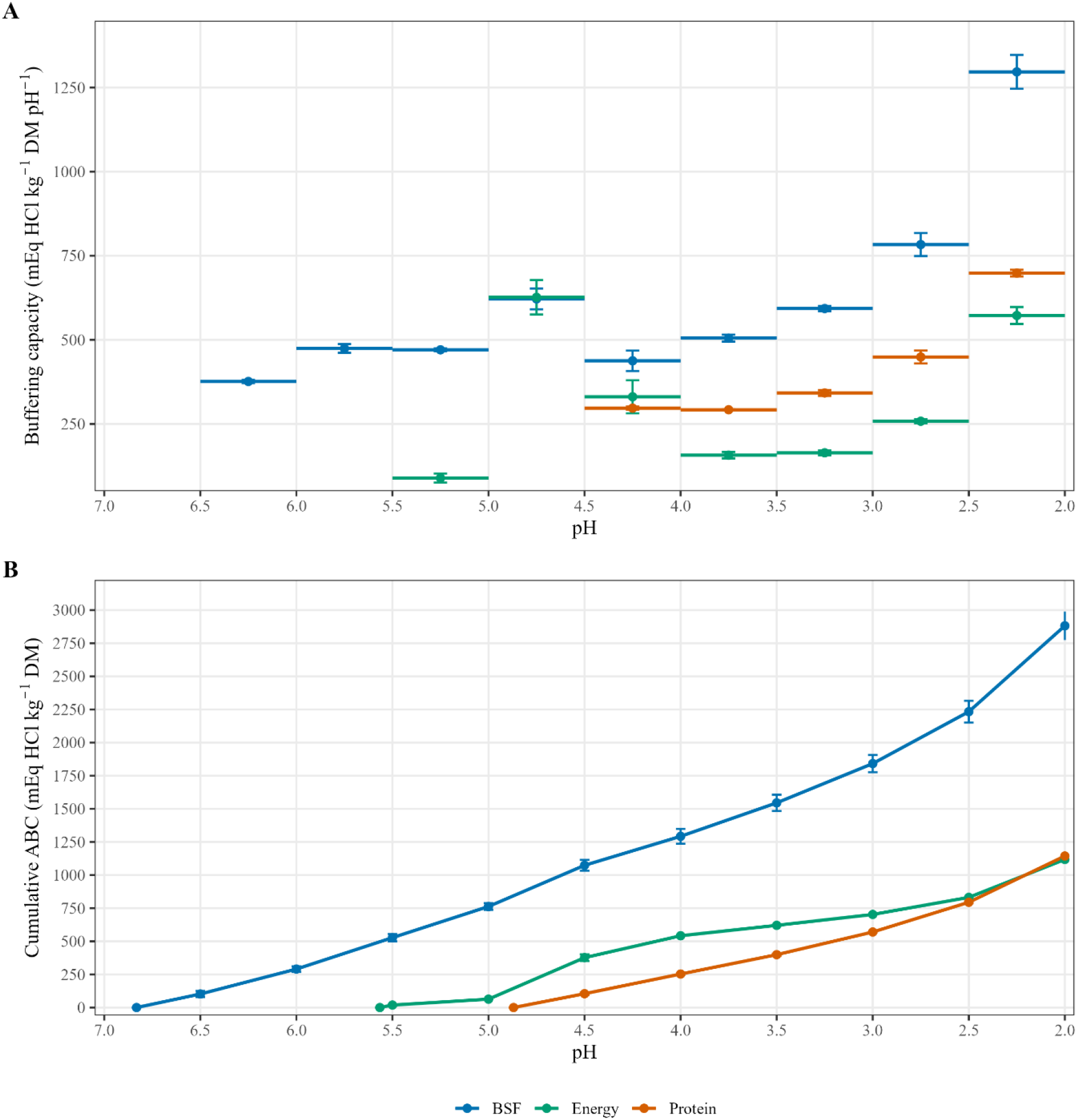
Buffering and acid-binding characteristics of black soldier fly (BSF), Protein core, and Energy core. (A) Buffering capacity, expressed as mEq HCl kg⁻¹ DM pH⁻¹, across complete 0.5-pH-unit intervals. Horizontal segments indicate the pH interval to which each value applies; points and error bars are positioned at the interval midpoint. (B) Cumulative acid-binding capacity (ABC), expressed as mEq HCl kg⁻¹ dry matter (DM), as a function of pH. Values are presented as means ± technical standard deviations (n = 2). The initial partial pH interval was excluded from panel A.

The pH-stat measurements of degrees of hydrolysis (Figure 2) showed that the Energy Core had the lowest degree of hydrolysis after 1 h, both at the low pepsin dose, with a value of 2.7 ± 0.1%, and at the high pepsin dose, with a value of 6.1 ± 0.1%. The same trend was observed after 6 h at the low dose and after 15 h at the high dose, with respective values of 6.0 ± 0.3% and 12.5 ± 0.1%.

**Figure 2.**
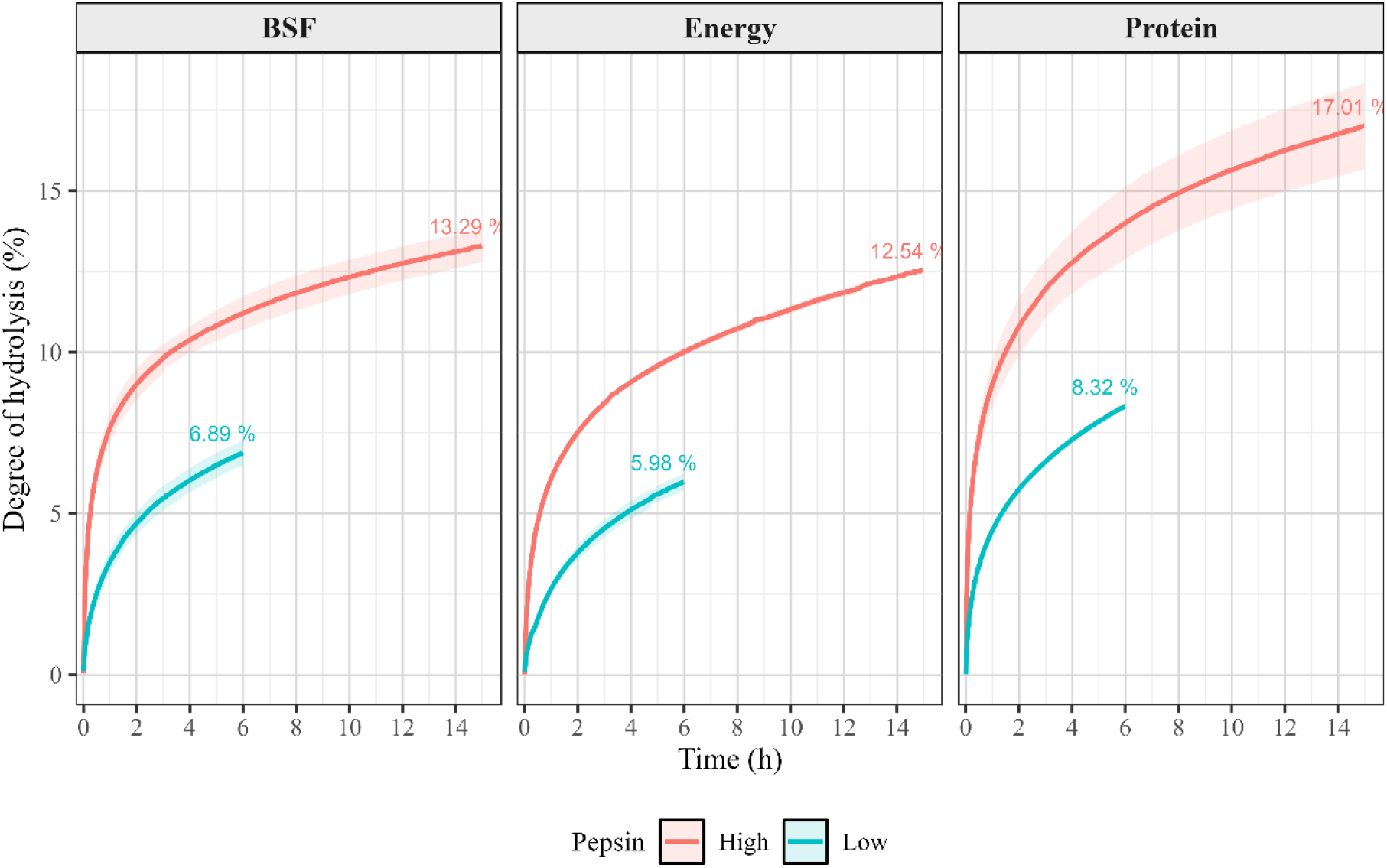
Degree of hydrolysis (DH, %) as a function of time (h) for Black Soldier Fly (BSF), Energy Core, and Protein Core measured by the pH-stat method. High and low pepsin dose treatments corresponded to 50 mg/mL (100 µL added; 5 mg pepsin) and 4 mg/mL (125 µL added; 0.5 mg pepsin) porcine pepsin, respectively. Lines represent the mean of two independent digestions and shaded areas represent ± SD (n = 2).

After 1 h, the Protein Core and BSF showed degrees of hydrolysis of 4.5 ± 0.1% and 3.5 ± 0.4%, respectively, at the low pepsin dose, and 9.0 ± 0.7% and 7.7 ± 0.5%, respectively, at the high dose. At the low pepsin dose after 6 h, the Protein Core and BSF reached values of 8.3 ± 0.0% and 6.9 ± 0.4%, respectively. Finally, at the high pepsin dose after 15 h, their degrees of hydrolysis were 17.0 ± 1.4% and 13.3 ± 0.5%, respectively.

## Discussion

This study compared the protein quality and degradation degrees of hydrolysis of a conventional organic protein core and a feed-grade BSF meal. Since protein quality depends not only on crude protein concentration but also on amino acid composition and availability, the sum of total amino acids of both ingredients was first determined. Our results showed that the protein core and BSF meal contained similar amounts of amino acids, representing approximately 50% of dry matter. In monogastric nutrition, particularly when insect-derived ingredients are included at substantial levels, relying solely on total nitrogen or on a non-specific nitrogen-to-protein conversion factor may lead to an overestimation of the actual protein content (Janssen et al., 2017).

The discrepancies between protein contents estimated from nitrogen conversion factors and those derived from amino acid analysis can be attributed to the presence of non-protein nitrogen in the feed ingredients. These differences were particularly pronounced in the BSF meal and are most likely related to the presence of chitin (Eggink & Dalsgaard, 2023; Soetemans et al., 2020). To further investigate this hypothesis, glucosamine, a chitin-derived amino sugar detected during amino acid analysis, was considered as an indicator of chitin-derived compounds. However, glucosamine results should be interpreted with caution and do not allow for a precise quantification of chitin in our samples. The analytical procedure used in this study was designed for amino acid determination and allows the detection of glucosamine, but it was not specifically developed or validated for chitin quantification (Luparelli et al., 2023; Smets C Van Der Borght, 2021). Therefore, these results mainly provide insight into the potential forms of non-protein nitrogen present in the tested ingredients.

Nevertheless, nitrogen-to-protein conversion factors only provide an estimate of protein content and cannot replace the determination of true protein or amino acids profile when accurate diet formulation is required. Unlike nitrogen-based estimations, true protein determination reflects the fraction of nitrogen associated with amino acids. It therefore provides a more reliable basis for evaluating the protein value of feed ingredients. Regarding amino acid composition, both the concentrations and the relative proportions observed in the present study were generally consistent with values reported in the literature (Eide et al., 2024; Huang et al., 2019; Liu et al., 2017). Nevertheless, the amino acid composition of BSF meals may vary depending on the rearing substrate, whereas the degree of defatting directly influences amino acid concentrations when expressed on a dry matter basis (Biasato et al., 2019; Crosbie et al., 2021).

The protein core and BSF meal exhibited relatively similar amino acid profiles for most indispensable amino acids (IAAs). This was also accompanied by only moderate differences in the profile of non-indispensable amino acids (NIAAs). Although the balance between IAAs and NIAAs may influence protein quality, the zootechnical relevance of NIAAs and the potential benefits of their supplementation remain poorly documented (Correia et al., 2023; Hou & Wu, 2017). Another notable difference between the two ingredients was their free amino acid (FAA) content. The BSF meal contained approximately 10% FAAs, corresponding to a concentration nearly twenty times higher than that of the protein core. Finally, amino acid composition alone does not provide information on amino acid digestibility or bioavailability. Black soldier fly proteins have previously been shown to exhibit amino acid digestibility comparable to that of conventional protein sources when fed to growing pigs (Crosbie et al., 2020).

In this study, we aimed to evaluate the availability of reactive lysine in the tested feed ingredients. As lysine is generally the first limiting amino acid and is particularly susceptible to heat damage (Brestenský et al., 2014), the thermal treatments applied to BSF proteins to comply with the European regulations governing processed animal proteins were considered a potential cause of reduced amino acid availability.

Our initial results indicated an apparent lysine damage of 18.4% in the BSF meal, compared with only 1.4% in the protein core. Additional experiments showed that the conversion of lysine to homoarginine depends strongly on the chemical form of lysine, with conversion yields of approximately 100% when lysine is protein-bound, 82% as a Lys-Lys dipeptide, and only 27% as free lysine. Consequently, failing to account for free amino acids, particularly in ingredients containing substantial amounts of them, leads to an overestimation of lysine damage. After correcting for the presence of FAAs, the estimated lysine damage in the BSF meal decreased to approximately 10.8%. The lower conversion yield observed for free L-lysine indicates that OMIU-mediated guanidination does not proceed with equal efficiency across all lysine forms. Consequently, reactive lysine may be underestimated in ingredients containing substantial amounts of free amino acids. The literature reported incomplete conversion of free L-lysine to homoarginine during OMIU-mediated guanidination (Hulshof et al., 2017). The present results extend these observations by showing that conversion is also incomplete when lysine is present as a Lys-Lys dipeptide. However, the conversion efficiency is considerably higher than for free L-lysine. This finding is particularly relevant for BSF meal because of its measurable free lysine content, leading to an overestimation of apparent lysine damage when no correction is applied.

Part of this apparent lysine damage may be explained by the Maillard reaction markers quantified in the present study. Furosine (FUR), an early marker of the Maillard reaction (Nielsen et al., 2022), was present at concentrations approximately five times higher in the BSF meal than in the protein core. Likewise, the advanced lysine-derived Maillard markers (Wei et al., 2018) Nε-carboxymethyllysine (CML) and Nε-carboxyethyllysine (CEL) were detected at concentrations approximately eight times higher in the BSF meal. Although methylglyoxal-derived hydroimidazolone 1 (MG-H1) is formed from arginine rather than lysine, it is also considered a marker of heat-induced protein modification (Qin et al., 2022), and in contrast to CML and CEL, the protein core contained approximately 20% more MG-H1 than the BSF meal, suggesting that these two matrices may respond differently to thermal processing.

However, thermal treatments applied to vegetable protein sources and processed animal proteins differ substantially. The heat treatments required for processed animal proteins are generally more severe and are therefore expected to induce greater lysine damage. Although FUR and advanced glycation end products (AGEs) were present at higher concentrations in the BSF meal, lysine damage remains comparable to that reported for other processed animal proteins, such as fish meal (Boucher et al., 2009). Nevertheless, the presence of Maillard reaction products raises questions regarding their potential effects on animal health (Teodorowicz et al., 2018; Zhang et al., 2020). Furthermore, extensive lysine damage may reduce the theoretical nutritional value of a feed ingredient, although this could potentially be compensated for by supplementation with synthetic L-lysine.

Another important aspect of this study was evaluating the buffering capacity of the tested ingredients, together with their protein degrees of hydrolysis, using the pH-stat method. Our results showed that the BSF meal exhibited a higher initial pH, cumulative acid-binding capacity, and buffering capacity than both the protein core and the energy core. Furthermore, under both high pepsin concentrations over an extended incubation period and lower pepsin concentrations over a shorter incubation period, the protein core consistently showed greater protein degradability than either the BSF meal or the energy core.

These results suggest that BSF meal requires a larger amount of hydrochloric acid to reach the acidic conditions required for optimal pepsin activation (Deng et al., 2021; Wang et al., 2023). Moreover, even once the optimal pH is reached, the proteins appear to be degraded less efficiently than those of the protein core. This observation may be particularly relevant for young animals such as post-weaning piglets, whose gastric acid secretion is not yet as efficient as that of adult pigs (Ferronato & Prandini, 2020). Consequently, high inclusion levels of black soldier fly meal may slow gastric protein digestion or reduce digestive efficiency. Together with its lower protein degradability, this could partly explain the reduction in animal performance reported in some studies when high levels of black soldier fly meal replace soybean-rich protein concentrates similar to those used in the present study.

The higher cumulative acid-binding capacity and buffering capacity of the BSF meal may partly be explained by its higher concentrations of certain minerals compared with the protein core. However, differences in mineral composition alone are unlikely to fully explain the observed differences. The molecular size of peptides and the presence of chitin in its native, non-purified form may also contribute to the lower degradability of BSF proteins and deserve further investigation (DiGiacomo & Leury, 2019; Liu et al., 2023).

Despite these observations, dietary acidification could represent a potential strategy to improve the digestive characteristics of BSF meal. Organic acids, such as citric acid (Nguyen et al., 2020), may help lower gastric pH and partly counteract the relatively high cumulative acid-binding and buffering capacities observed in the present study. However, this hypothesis would require experimental confirmation under in vivo conditions.

It would also be of interest to evaluate different energy cores to determine whether their composition could partly compensate for these buffering effects. Likewise, formulating the mineral premix according to the composition of the final diet containing BSF meal, rather than solely based on the energy core, could represent another approach deserving further investigation.

## Conclusion

This study assessed the protein quality and protein degradability of a feed-grade defatted black soldier fly meal and a conventional protein core intended for post-weaning piglets. Defatted black soldier fly meal exhibited a protein content and amino acid profile comparable to those of the protein core. However, it also contained substantially higher levels of free amino acids and Maillard reaction products, resulting in an apparent overestimation of lysine damage when reactive lysine was determined by OMIU-mediated guanidination without accounting for free lysine. In addition, BSF meal exhibited a higher buffering capacity and cumulative acid-binding capacity together with a lower protein degradability than the protein core. These characteristics may represent important constraints when formulating diets for young pigs with immature gastric function. Overall, the present findings contribute to a better nutritional characterization of BSF meal and highlight several avenues for improving its use in post-weaning piglet diets, including a more accurate assessment of reactive lysine, optimization of dietary acidification, and formulation strategies accounting for the buffering properties of the complete diet. This research provides new insights into the proteins of black soldier flies and identifies new factors that are important to consider when formulating diets containing this feed.

## Acknowledgements

The authors would like to thank the Walloon Region (Service Public de Wallonie; DGO6) for funding Hugo Luttenschlager through the ASTIPPOR project (D65-1438), obtained under the Walloon Recovery Plan (https://www.wallonie.be/en/plans-wallons/plan-de-relance-de-la-wallonie), which enabled the production of the insect proteins used in this study. The authors also thank Wallonie-Bruxelles International for awarding a WBI Excellence Fellowship to Hugo Luttenschlager, allowing him to conduct his analyses at the Animal Science (Nutrition Group) laboratory at Wageningen University & Research. The authors would like to thank Extratex S.F.I. (Pont-Saint-Vincent, France) for performing the supercritical CO₂ extraction.

## Author’s contribution

**Hugo Luttenschlager**: Conceptualization, Sample preparation and production, Laboratory analyses, Data analyses, Writing – Original draft, Funding acquisition. **Niels Wever**: Conceptualization, Sample preparation and production, Laboratory analyses, Data analyses, Writing – review & editing. **Wouter Hendriks**: Conceptualization, Supervision, Writing – review & editing. **Yves Beckers**: Conceptualization, Writing – review & editing, Funding acquisition. **Samuel Jansseune**: Laboratory analyses, Data analyses, Writing – review & editing. **Joachim Carpentier**: Sample preparation and production, Writing – review & editing. **Aurore Richel**: Technical support, Writing – review & editing. **Frédéric Francis**: Technical support, Writing – review & editing. **Rudy Caparros Megido**: Conceptualization, Supervision, Original draft writing, Funding acquisition.

## Data availability

Data can be requested directly from the reference author (Hugo Luttenschlager) at and from the supervisor (Rudy Caparros Megido) at.

