## Supplementary material for "Assessing protein quality, reactive lysine and gastric digestion characteristics of feed-grade black soldier fly meal: Implications for post-weaning piglet feed formulation": Table S1

Table S1: MRM transitions and their optimal conditions as used for this experiment.

| Range | Name | Precursor | Fragment | Cone voltage | Collision Energy |
|---|---|---|---|---|---|
| [min] | | [m/z] | [m/z] | [V] | [eV] |
| 2.5-4.5 | MG-H1 | 229 | 166 | 26 | 16 |
| 2.5-4.5 | MG-H1-d4 | 233 | 170 | 26 | 16 |
| 3.0-5.0 | Furosine | 255 | 130 | 26 | 14 |
| 3.0-5.0 | Furosine-d4 | 259 | 134 | 26 | 14 |
| 3.5-5.5 | CEL | 219 | 130 | 28 | 12 |
| 3.5-5.5 | CEL-d4 | 223 | 134 | 28 | 12 |
| 3.5-5.5 | CML | 205 | 130 | 28 | 12 |
| 3.5-5.5 | CML-d4 | 209 | 134 | 28 | 12 |
